# High-dose Vitamin C Blocks HOCl Production by Myeloperoxidase: A Potential Therapeutic Strategy

**DOI:** 10.1101/2024.12.05.627077

**Authors:** Alexander V Peskin, Nicholas J Magon, Stephanie M Bozonet

## Abstract

High-dose vitamin C therapy for cancer, originally advocated by Linus Pauling (Proc Natl Acad Sci, 1976, 73, 3685–3689), remains a subject of ongoing debate. In this study, we investigate why only pharmacological doses are effective and explore the reasons behind inconsistent therapeutic outcomes. Our data suggest that the bona fide cause of toxicity was oxidized vitamin C rather than hydrogen peroxide. We found that vitamin C at millimolar concentrations, directly inhibits hypochlorous acid generation by myeloperoxidase, through competition with chloride rather than by scavenging the hypochlorous acid that is formed. Products of vitamin C oxidation reacted with the thiols of peroxiredoxin 2 and GAPDH, but failed to react with the cysteine of p16^INK4a^. The growth and viability of Jurkat cells were affected by oxidized vitamin C. These experiments were conducted in the presence of catalase, demonstrating that the biological effects were due to the products of vitamin C oxidation and not hydrogen peroxide. These findings may have practical implications for the treatment of cancer and diseases in which the deleterious effects of neutrophil activation are observed. For intravenous administration of pharmacological vitamin C to have a beneficial effect, its concentration in the blood must be maintained at millimolar levels and this can only be achieved via maintenance infusion. As a proof-of-concept, our data suggest that to enhance anticancer therapy interventions, it is crucial to implement treatments that facilitate the oxidation of vitamin C in the bloodstream.

> ***Progress in the art of healing has always been induced by progress in basic knowledge.***

> ***Albert Szent-Gyorgyi (Woods Hole, Mass., USA, 1973)***

**Graphical abstract:** 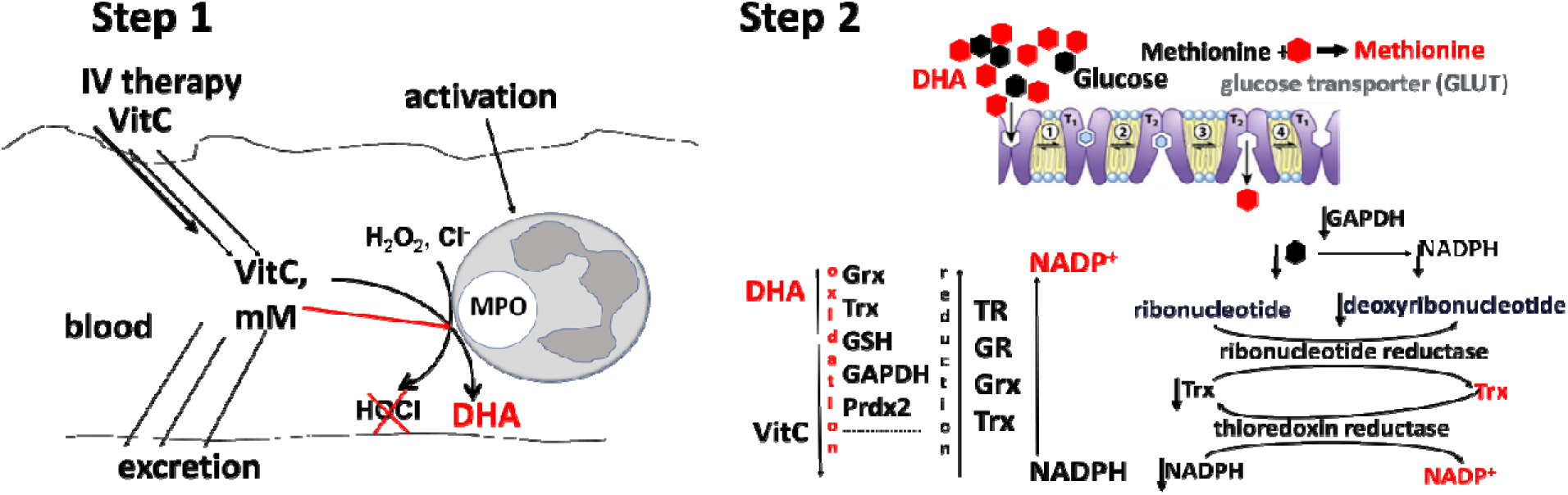

## INTRODUCTION

The seminal publication by Linus Pauling in 1976, on the effects of high-dose vitamin C (ascorbate) in cancer (1), triggered investigations aimed at understanding the biological role of vitamin C in humans. In addition to being a potent free radical scavenger, it has been shown that ascorbate is a cofactor for a family of biosynthetic and gene-regulatory monooxygenase and dioxygenase enzymes and, as such, is involved in the synthesis of collagen, carnitine, and hormones (2,3). Members of this enzyme family include the 2-oxoglutarate-dependent dioxygenases which regulate their targets through hydroxylation. In the context of cancer, these enzymes downregulate hypoxia inducible factors, key drivers of the hypoxic response and tumour cell survival. They also regulate histone and DNA demethylases, such as the ten-eleven translocase and Jumonji-C-domain-containing proteins, which are involved in the epigenetic control of oncogenes and tumour suppressors (4–6).

Despite significant advances in our understanding of the biological effects of vitamin C, there is still no answer to the question of why pharmacological levels (achieved with higher-than-normal doses for therapeutic or medical purposes) of vitamin C are necessary to observe anticancer effects.

Currently, the most favored explanation attributes the anticancer effects to hydrogen peroxide (H_2_O_2_) formed by metal-catalyzed oxidation of high levels of ascorbate (7,8). However, doubts about the feasibility of this explanation have been raised (9,10).

We hypothesized that the anticancer effect of high-dose vitamin C therapy results from its oxidation products. Dehydroascorbate (DHA), the immediate product of the two-electron oxidation of vitamin C, is known to oxidize thiols (11–13) and is toxic to cancer cells (14–17). However, due to its instability, DHA cannot be used clinically (18,19). The observed toxic effect of high-dose vitamin C has been attributed to H_2_O_2_ accumulation from vitamin C autoxidation. However, H_2_O_2_ can further react with vitamin C to produce DHA. If H_2_O_2_ were the primary driver of toxicity, removing it would prevent DHA formation and, consequently, the cytotoxicity of high-dose vitamin C. Yet, if DHA is the true toxic agent, then adding pre-oxidized vitamin C together with catalase should still kill cancer cells in culture. In blood, the autoxidation of high-dose vitamin C cannot sustain sufficient H_2_O_2_ levels to generate high DHA concentrations due to H_2_O_2_ diffusion into erythrocytes packed with peroxide-removing enzymes.

Vitamin C can be effectively oxidized in blood by myeloperoxidase (MPO), heme-containing peroxidase most abundantly expressed by white blood cells (WBC). Activated MPO uses H_2_O_2_ to oxidize Cl^-^ to produce hypochlorous acid (HOCl) which, as well as being bactericidal, can have unwanted side effects (20). HOCl reacts quickly with ascorbate, the immediate product of two-electron oxidation of ascorbate is DHA:

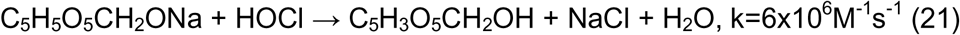

Activated MPO can also oxidize ascorbate directly (22). In its basal state MPO reacts with H_2_O_2_ and is converted to Compound I. Compound I then reacts with Cl⁻ at a second-order rate constant of ∼10□ M⁻¹s⁻¹, producing HOCl and returning to its basal state (23). Compound I may alternatively react with ascorbate to form Compound II, which reacts with another molecule of ascorbate before converting back to MPO in its basal state. In comparison to Cl^−^, the second-order rate constant for the reaction of ascorbate with Compound I is much faster, ∼10^6^ M^−1^s^−1^ (22). But 100 μM vitamin C, which is upper level of normal physiological concentration in blood, cannot prevent HOCl generation by MPO (24). We suggest that pharmacological levels of ascorbate, which can exceed 10 mM in the blood (25), would outcompete physiological concentrations of Cl^−^, and therefore MPO would produce oxidized ascorbate rather than HOCl under these conditions.

Recent publications highlight the targeting of thiol metabolism as a promising approach to anti-cancer therapy (26, 27) and these are supported by the findings that peroxiredoxins and thiol metabolism are upregulated in tumours (28,29). Peroxiredoxins are thiol-containing proteins that cycle between oxidized and reduced forms, and a growing body of evidence indicates their critical role in cell signalling (30–32). It has been reported that inhibition of peroxiredoxins results in leukemic cell differentiation (33). Peroxiredoxin 2 (Prdx2) is essential for proliferation and S-phase progression, and it promotes colorectal cancer cell-cycle progression through activation of the p38 MAPK pathway (34). Thiol oxidation in glyceraldehyde-3-phosphate dehydrogenase (GAPDH) inhibits its enzymatic activity (35), which is essential for glucose metabolism, and tumour growth in vivo is suppressed when the GAPDH redox switch is disabled in tumour cells (36).

In this paper, we assessed whether DHA can inhibit Prdx2 and GAPDH, both recognized targets for anticancer treatment.

Protein p16^INK4a^ is a negative regulator of cell division, inhibiting the cyclin dependent kinases 4 and 6 and preventing progression of the cell cycle. The loss of p16^INK4a^ activity is widely believed to be a common and important event in the development of cancer. Oxidation of p16^INK4a^ leads to the formation of disulfide-bridged dimers that subsequently form amyloid fibrils. While amyloid fibrils typically form spontaneously, p16^INK4a^ is the first example of strictly oxidation-induced amyloid formation (37).

Assessing the oxidation of p16^INK4a^ by DHA would help determine whether high doses of oxidized vitamin C could have pro-carcinogenic potential.

The anticancer effects of vitamin C and DHA were assessed on Jurkat cells, a leukemic human T lymphocyte cell line commonly used for cancer studies.

## METHODS

### Proteins

Recombinant human Prdx2 was prepared as described (38) and provided by Paul Pace. Recombinant human p16^INK4a^ was a gift from Christoph Göbl (37). Human myeloperoxidase was from Tony Kettle (39). Rabbit muscle GAPDH, bovine catalase and other chemicals were from Sigma-Aldrich. Catalase protein concentration was estimated spectrophotometrically (□_405_ = 120 mm^−1^ cm^−1^), and activity was measured by following the loss of added H_2_O_2_ at 240 nm as advised by Sigma. From these data we calculated a specific activity of 4700 units/mg.

All thiol-containing proteins were reduced by the addition of 10 mM dithiothreitol (DTT) for 1 h immediately before use. Excess DTT was removed using Micro Bio-Spin 6 columns (Bio-Rad), which were prewashed with deionized water and then with 5 ml of 20 mM phosphate buffer, pH 7.4, containing 0.1 mM diethylenetriamine-penta-acetic acid (DTPA) and purged with argon. Experiments were conducted in 20 mM sodium phosphate buffer, pH 7.4, containing 0.1 mM DTPA.

### Sources of oxidized ascorbate

1. Commercial dehydroascorbic acid (Sigma-Aldrich) was dissolved immediately before use in 20 mM phosphate buffer, containing 0.1 mM DTPA, adjusted to pH 7.4, then mixed with catalase, 10 µg/ml.
2. Sodium ascorbate was dissolved in deionized water immediately before use. Hypochlorous acid (HOCl) was purchased as commercial chlorine bleach from Household and Body Care (Auckland, NZ) The stock’s concentration was determined spectrophotometrically using ε _292_ = 350 M^−1^ cm^−1^ at pH 12. HOCl from stock was diluted in 20 mM phosphate buffer and added to ascorbate dropwise, during vigorous mixing at a molar ratio of 1:2. Stoichiometry of the reaction is 1:1 (21).
3. Myeloperoxidase from stock was initially diluted in H_2_O and further diluted for use (to 20 or 40 nM) in phosphate buffer (20 mM, pH 7.4 containing 0.1 mM DTPA) with varying concentrations of NaCl and/or ascorbate. After mixing, H_2_O_2_ was added for 5 min, then catalase (10 µg/ml) was added. For analysis of methionine oxidation, protein was removed using 3 kDa centrifuge filters (Millipore) before analysis by mass spectroscopy.

### GAPDH activity assay

The activity of GAPDH, 50 μg/ml either untreated or treated with oxidized ascorbate, was measured, in Tris buffer (Tris-HCl, 50 mM, pH 8) containing EDTA (0.2 mM), glyceraldehyde 3-phosphate (0.4 mM), NAD^+^ (1 mM) and sodium arsenate (15 mM), by monitoring the initial reduction rate of NAD^+^ as an increase in absorbance (A_340nm)_ (40) using a SpectraMax iD3 plate reader with SoftMax Pro 7.1 software (Molecular Devices, San Jose, California, USA) as described earlier (35).

### Cell culture conditions and treatments

Jurkat cells (Clone E6-1, TIB-152 from ATCC) were cultured in Roswell Park Memorial Institute (RPMI) 1640 medium with 1% penicillin/streptomycin (Gibco, from Thermo Fisher Scientific, Auckland, NZ) and 10 % foetal bovine serum (Moregate BioTech, Hamilton, New Zealand). Cells were seeded in 24-well culture dishes, at a density of 0.25 × 10^6^/ml, and, where appropriate, catalase (20 µg/ml) was added to the culture medium before treatment. Ascorbate was oxidized by reaction with HOCl immediately before addition to the cells. Concentrations of ascorbate were chosen to inflict visible effect on cells. Phase contrast images were captured after 24 hours with an Olympus DP21 camera using a 20x magnification objective, then the cells were counted and viability measured: propidium iodide (PI) was added and the cells analysed using a Cytoflex S flow cytometer (Beckman Coulter).

### Mass spectrometry

For whole protein analysis, samples containing 1 µg of protein were injected onto an Accucore-150-C4 (50 x 2.1-mm, 2.6-µm) column (60 °C) using a Dionex Ultimate 3000 HPLC system coupled to a Velos Pro mass spectrometer (Thermo Scientific, San Jose, CA). Proteins were eluted with an acetonitrile gradient from 90% solvent A (0.1% formic acid in water) and 10% solvent B (0.1% formic acid in acetonitrile) to 80% solvent B over 4.6 min at a flow rate of 400 µl/min. Mass spectra were acquired between m/z 400 and 2000 in positive ion mode. The spectra of each protein peak were averaged using Thermo Xcalibur Qual Browser 4.2.47 (Thermo Fisher Scientific) and deconvoluted to yield the molecular masses and relative intensities using ProMass for Xcalibur (version 2.8 rev 5; Novatia LLC, Monmouth Junction, NJ).

For detection of methionine and its oxidation products a multiple reaction monitoring (MRM) method was set up (Table 1), using a Sciex 6500 QTrap mass spectrometer (Framingham, MA, USA) coupled to an Infinity 1290 LC system (Agilent, Santa Clara, CA, USA). Samples were stored on the autosampler tray at 5 °C prior to injection. An Imtakt Intrada Amino Acid column (150 x 3.0 mm) was used for chromatographic separation using acetonitrile containing 0.1% formic acid (Solvent A) and 100 mM ammonium formate (Solvent B). The column temperature was set to 40 °C. The column was equilibrated with 86% Solvent A and 14% Solvent B for 6 minutes and then a linear gradient was run for 14 minutes to 100% Solvent B to achieve separation. The column was then flushed with 100% Solvent B for 2 minutes and re-equilibrated at initial conditions for 4 minutes. A flow rate of 0.25 mL/minute was used. Data were analyzed using Analyst 1.7.2 (Sciex, Framingham, MA, USA).

**Table 1.**
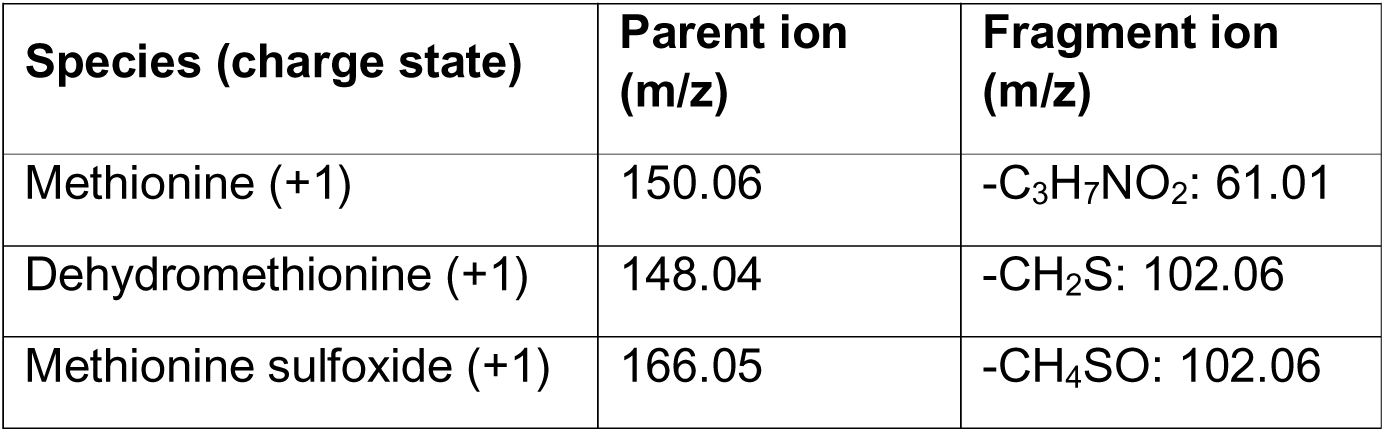
The m/z values for the parent and fragment ions that were used to quantify methionine species by LC/MS.

## RESULTS

### 1. Inhibition of HOCl generation by myeloperoxidase in the presence of pharmacological levels of ascorbate

Measuring the inhibition of HOCl generation by MPO in the presence of ascorbate is complicated by the very fast reaction of HOCl with ascorbate, which has a second order rate constant of 6×10^6^ M^−1^s^−1^(21). It is therefore necessary to distinguish between the genuine inhibition of HOCl generation by MPO, and scavenging of the formed HOCl by ascorbate. To address this issue, we introduced methionine to the reaction mixture: the reaction of methionine with HOCl has a second order rate constant of 3.8×10^7^ M^−1^s^−1^ (41) and therefore, at equimolar with ascorbate concentrations, methionine should scavenge most of the HOCl.

We used purified MPO and mass spectrometry to analyse methionine oxidation by MPO-generated HOCl, in the presence and absence of ascorbate.

Hydrogen peroxide oxidizes methionine to form methionine sulfoxide while HOCl oxidizes methionine to a mixture of dehydromethionine (DHM) and methionine sulfoxide (42), both of which can be measured using mass spectrometry. We measured the intensity of signals corresponding to methionine-related species, generated by MPO, when it was incubated with methionine in the presence of 100 mM NaCl after the addition of H_2_O_2_. Fig. 1 shows chromatograms of the species detected in the model mixture of methionine with HOCl, and these include methionine, two isomers of dehydromethionine, and methionine sulfoxide. With MPO activated by H_2_O_2_ in the presence of NaCl we observed the formation of DHM and methionine sulfoxide, which were decreased by half when ascorbate was present (Fig. 2). The degree of inhibition of methionine oxidation was not affected by lowering the concentration of NaCl to 10 mM. The fact that inhibition of methionine oxidation was independent of the NaCl concentration may be explained by the direct oxidation of ascorbate by MPO and subsequent oxidation of methionine by oxidized ascorbate.

**Figure 1.**
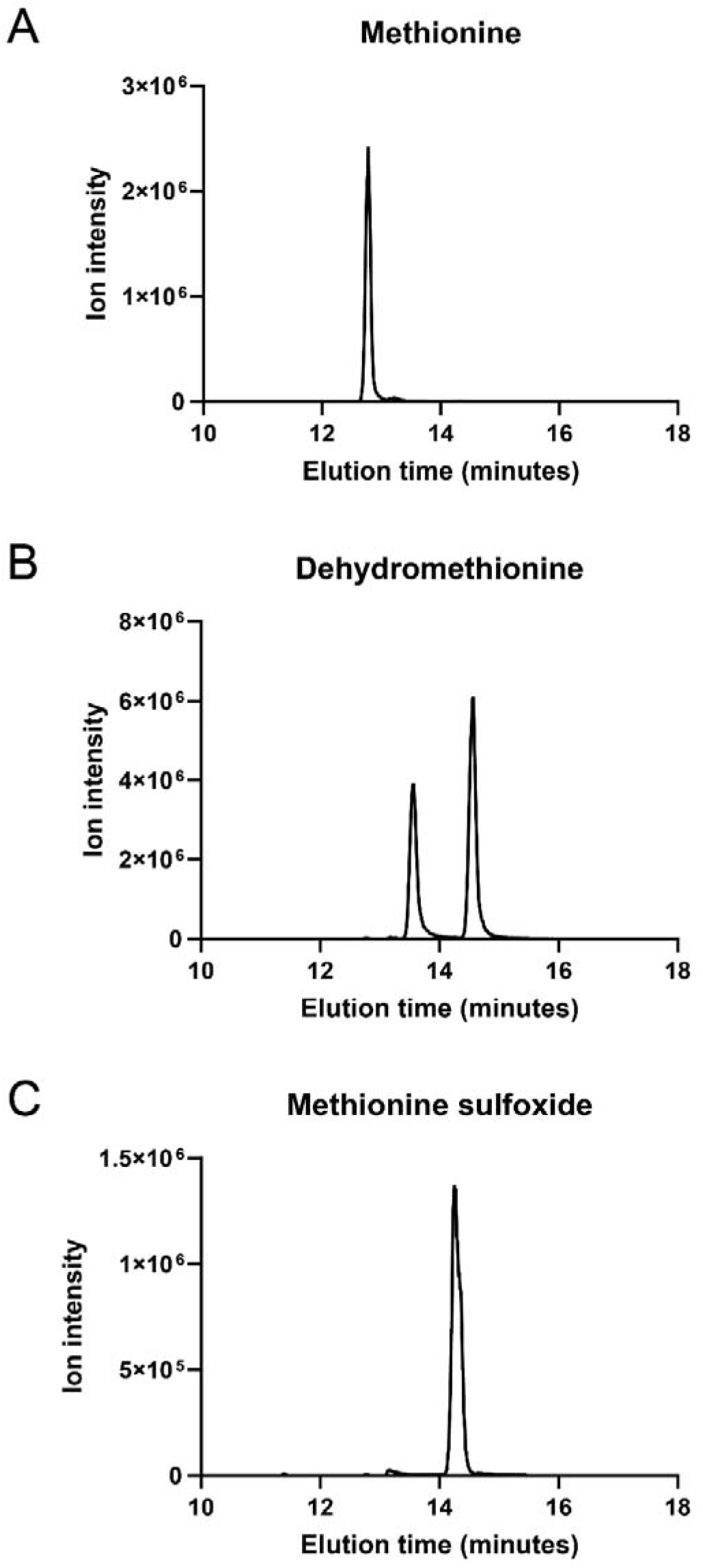
Chromatograms of methionine, dehydromethionine, and methionine sulfoxide. Methionine (10 mM) was mixed with HOCl (5 mM) in phosphate buffered saline, and 5 μL was injected for analysis.

**Figure 2.**
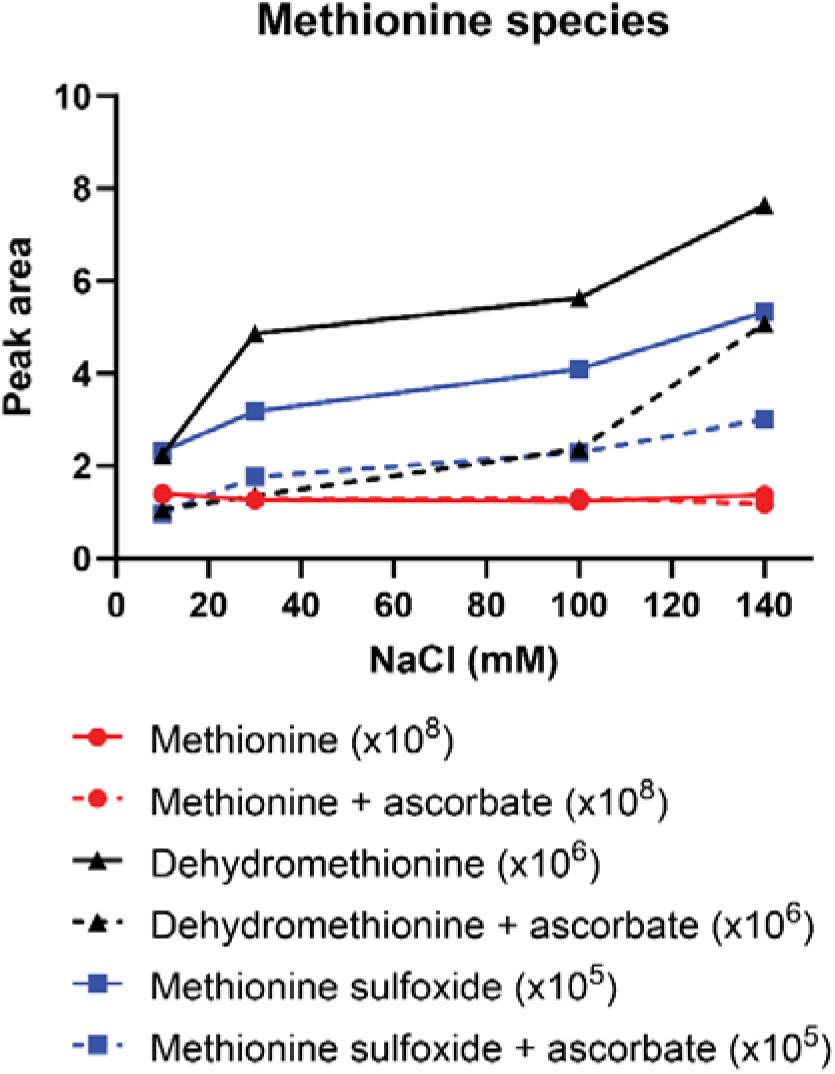
Inhibition of HOCl generation by myeloperoxidase in the presence of ascorbate. The reaction mix contained MPO (40 nM) and methionine (10 mM) and the reaction was started by the additon of H_2_O_2_ (40 µM). Where present, the ascorbate concentration was 10 mM and catalase (10 µg/ml) was added after 5 min. Then protein was removed from samples before analysis by mass spectrometry (see Methods). Recorded species: methionine (red), methionine sulfoxide (blue), dehydromethionine (black), in the absence (solid lines) and presence (dashed lines) of ascorbate. The intensity of the signals were ×10^8^ for methionine, ×10^5^ for methionine sulfoxide and ×10^6^ for dehydromethionine.

To assess whether direct oxidation of ascorbate by MPO leads to methionine oxidation, we conducted two sets of experiments. First, we delayed by 3 min the addition of methionine to the mixture containing MPO, NaCl, ascorbate and H_2_O_2_, to which catalase was added after 5 min. In this experimental setup, all of the HOCl would be scavenged by ascorbate before the addition methionine. However, in this case, we still observed the formation of DHM and methionine sulfoxide (Table 2). Second, we excluded NaCl from the mixture and looked for methionine oxidation at various concentrations of H_2_O_2_ (Table 3). Methionine oxidation occurred in the absence of NaCl (Table 3) and the formation of DHM showed a linear dependence on the concentration of H_2_O_2_ (Fig.3).

**Figure 3.**
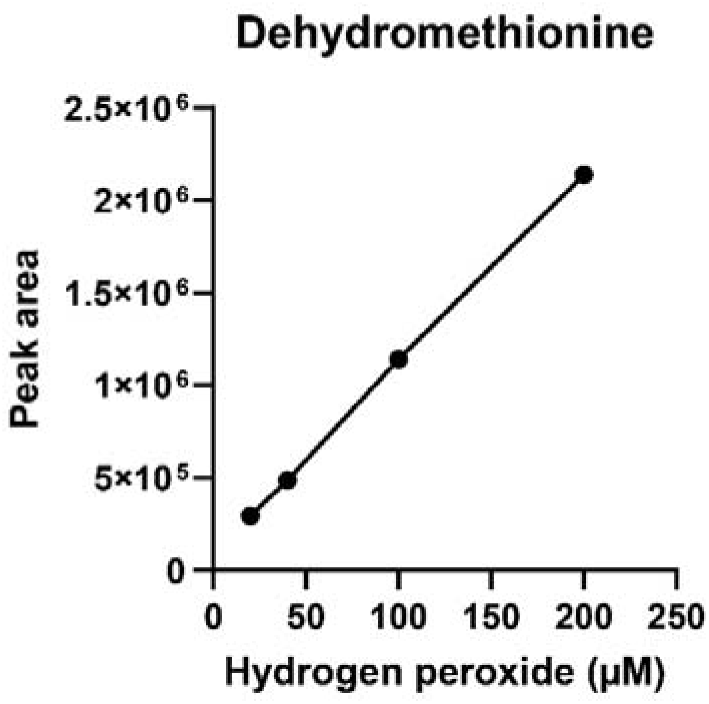
Dehydromethionine formation by MPO in the presence of ascorbate and absence of NaCl. The reaction mix contained MPO (40 nM) and ascorbate (10mM). After 5 min catalase (10 µg/ml) was added followed by methionine (10 mM). Then protein was removed from samples before analysis by mass spectrometry (see Methods).

**Table 2.** Oxidation of methionine by the products of ascorbate oxidation.

|  | Signal intensity |  |  |
| --- | --- | --- | --- |
|  | Methionine | Dehydromethionine | Methionine sulfoxide |
| Full system<br>100 mM NaCl | 1.64E+08 | 5.62E+06 | 4.32E+05 |
| Full system<br>minus<br>methionine<br>plus<br>ascorbate<br>100 mM NaCl<br>+ methionine<br>after 3 min | 1.56E+08 (95%) | 1.25E+06 (22%) | 2.33E+05(54%) |

Full system – MPO, methionine, H<sub>2</sub>O<sub>2</sub>. Each species was quantified by LC/MS using an MRM method (Table 1).

**Table 3.** Oxidation of methionine by MPO in reaction with H_2_O_2_ and ascorbate in the absence of NaCl. Each species was quantified by LC/MS using an MRM method (Table 1).

|  | Signal intensity |  |  |
| --- | --- | --- | --- |
|  | Methionine | Dehydromethionine | Methionine sulfoxide |
| MPO, Asc, Meth<br>+ 20 μM H<sub>2</sub>O<sub>2</sub> | 1.16E+08 | 3.38E+05 | 4.46E+05 |
| MPO, Asc, Meth<br>+ 40 μM H<sub>2</sub>O<sub>2</sub> | 1.11E+08 | 5.94E+05 | 4.92E+05 |
| MPO, Asc, Meth<br>+ 100 μM H<sub>2</sub>O<sub>2</sub> | 1.11E+08 | 1.20E+06 | 5.54E+05 |
| MPO, Asc, Meth<br>+ 200 μM H<sub>2</sub>O<sub>2</sub> | 1.16E+08 | 2.35E+06 | 4.95E+05 |

In contrast, the signal of methionine sulfoxide remained unchanged (Table 3), suggesting that it corresponds to the methionine sulfoxide present in the methionine stock solution.

We analysed commercially sourced methionine and methionine sulfoxide and, after equal loading, detected the following: the methionine, showed signal intensity of 4.57×10^7^ and 1.77×10^5^ for methionine and methionine sulfoxide, respectively; and in the methionine sulfoxide, these were 5.41×10^7^ and 1.78×10^5^ signals for methionine sulfoxide and methionine, respectively. In both chemicals the signal corresponding to DHM was in the range of 10^4^.

It should be noted that the reaction of Compound I with Cl^−^ is complex and dependent on the concentration of Cl^−^. Moreover, the efficacy of Compound I formation has a dependence on the amount of H_2_O_2_ available (23). Therefore, characterisation of the detailed mechanism behind the inhibition of HOCl generation by ascorbate is warranted.

Collectively, these data indicate that pharmacological levels of ascorbate can inhibit HOCl production by MPO and that ascorbate can react with MPO to produce species able to oxidize methionine. Little is known about the ability of the products of ascorbate oxidation to react with methionine (43). Further investigation into the mechanism of methionine oxidation by the products of ascorbate oxidation is warranted to determine whether DHM is a characteristic feature of the reaction.

### 2. Effects of oxidized ascorbate on thiol-containing proteins

#### 2.1. Peroxiredoxin 2

Reduced Prdx2 treated with oxidized ascorbate was analysed by LC/MS. Oxidized ascorbate was either in the form of commercial DHA or generated by MPO.

Prdx2 has two reactive cysteines and upon oxidation forms a covalent dimer with either one or two disulfide bonds. Prdx2 reacts rapidly with H_2_O_2_ and is oxidized by even trace amounts present in buffers (44). Catalase was therefore added to the oxidized ascorbate before mixing with Prdx2 to prevent oxidation of the Prdx2 cysteines by H_2_O_2_. Fig. 4 and Fig. 5 show that oxidized ascorbate resulted in the loss of monomeric Prdx2 and formation of Prdx2 dimer. We also observed species with masses corresponding to a parent dimer plus 39 - 42 Da, 106 Da and 208 - 214 Da (Figs. 4B and 5C, inserts).

**Figure 4.**
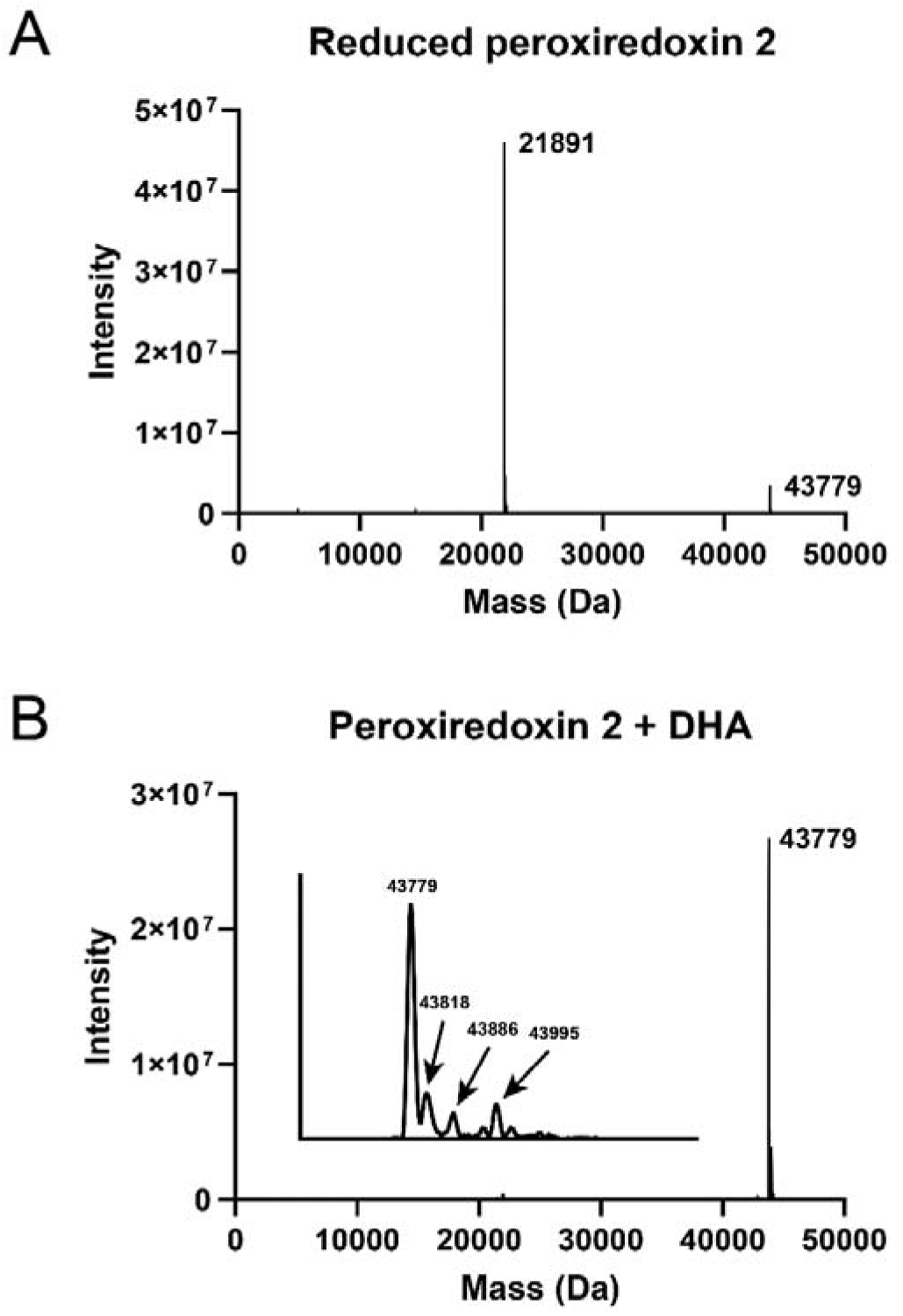
Oxidation of Prdx2 by DHA. LC/MS analysis of whole protein. Deconvoluted data are shown for reduced Prdx2, 5 µM (A) and Pdx2 after the mixture of DHA, 500 µM with catalase, 10 µg/ml was added to protein for 10 min incubation (B).

**Figure 5.**
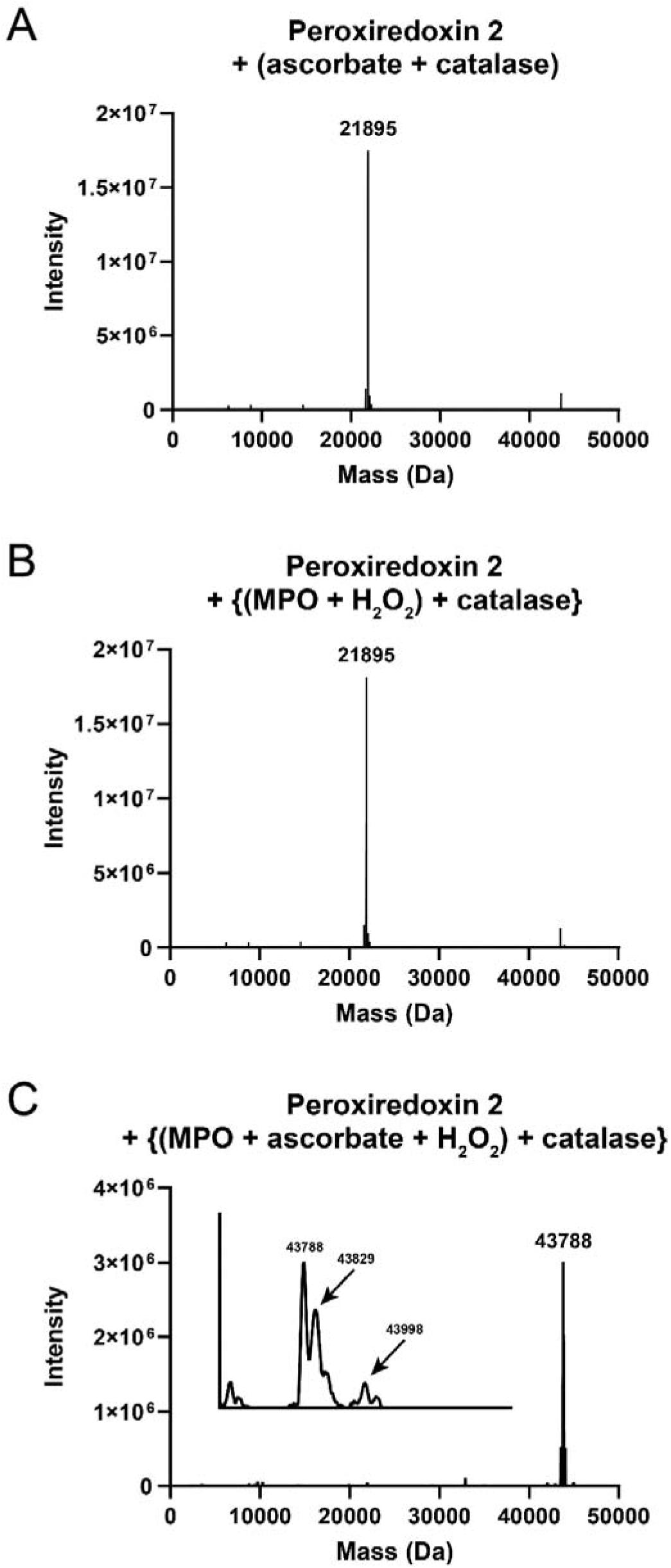
Oxidation of Prdx2 by the products of ascorbate oxidation pre-formed by HOCl. Whole protein was analysed by LC/MS. Prdx2, 5 µM; MPO, 20 nM; ascorbate, 200 µM; H_2_O_2_, 40 µM; catalase, 10 µg/ml. A - Prdx2 was added to the mixture of ascorbate with catalase; B - Prdx2 added to MPO, catalase was added after H_2_O_2_; C – H_2_O_2_ was added to the mixture of MPO with ascorbate, after 5 min catalase was added followed by Prdx2 and the mixture was incubated for an additional 10 minutes. 100 mM NaCl was present. Therefore, ascorbate was oxidized by HOCl as 200 µM ascorbate could not outcompete Cl^−^.

Adducts of glutaredoxin have been shown to form in the presence of oxidized ascorbate, with an increase in molecular weight of 112 Da resulting from the addition of the 5-carbon fragment of DHA to the protein thiol (13). In our experiments the accuracy of the deconvoluted whole protein masses was within 7 Da of the theoretical masses (45). We therefore suggest that the oxidized ascorbate may form adducts on Prdx2, similar to those observed for glutaredoxin: the addition of 106 Da may be an adduct of the 5-carbon fragment of DHA, and adducts with additional masses in the range of 208 - 214 Da may represent two adducts (each of 104 - 107 Da) on the two cysteines not involved in the disulfide bond of the Prdx2 dimer.

Formation of adducts with both Prdx2 cysteines has been shown for other electrophiles (46), and the addition of 39 - 42 Da could be attributed to an acetaldehyde adduct which may be produced from the decomposition of DHA (18, 19).

Variation in the adduct profiles formed with the two sources of oxidized ascorbate may result from differences in reactions of ascorbate oxidation used. Though regardless of the source, the oxidized ascorbate caused complete dimerization of Prdx2. Further investigation is warranted into these adducts, their stability and reversibility, and the impact of DHA decomposition on Prdx2.

#### 2.2. GAPDH

The enzymatic activity of GAPDH depends on the presence of reduced cysteine residues and oxidation of these results in the loss of activity (Fig. 6A). The addition of oxidized ascorbate to reduced GAPDH resulted in a concentration-dependent loss of enzymatic activity (Fig. 6B). The concentration-dependent inhibition of GAPDH activity by DHA was somewhat higher than we observed with H₂O₂ (35). But more data are needed for quantitative comparison in efficacy of these two oxidants.

**Figure 6.**
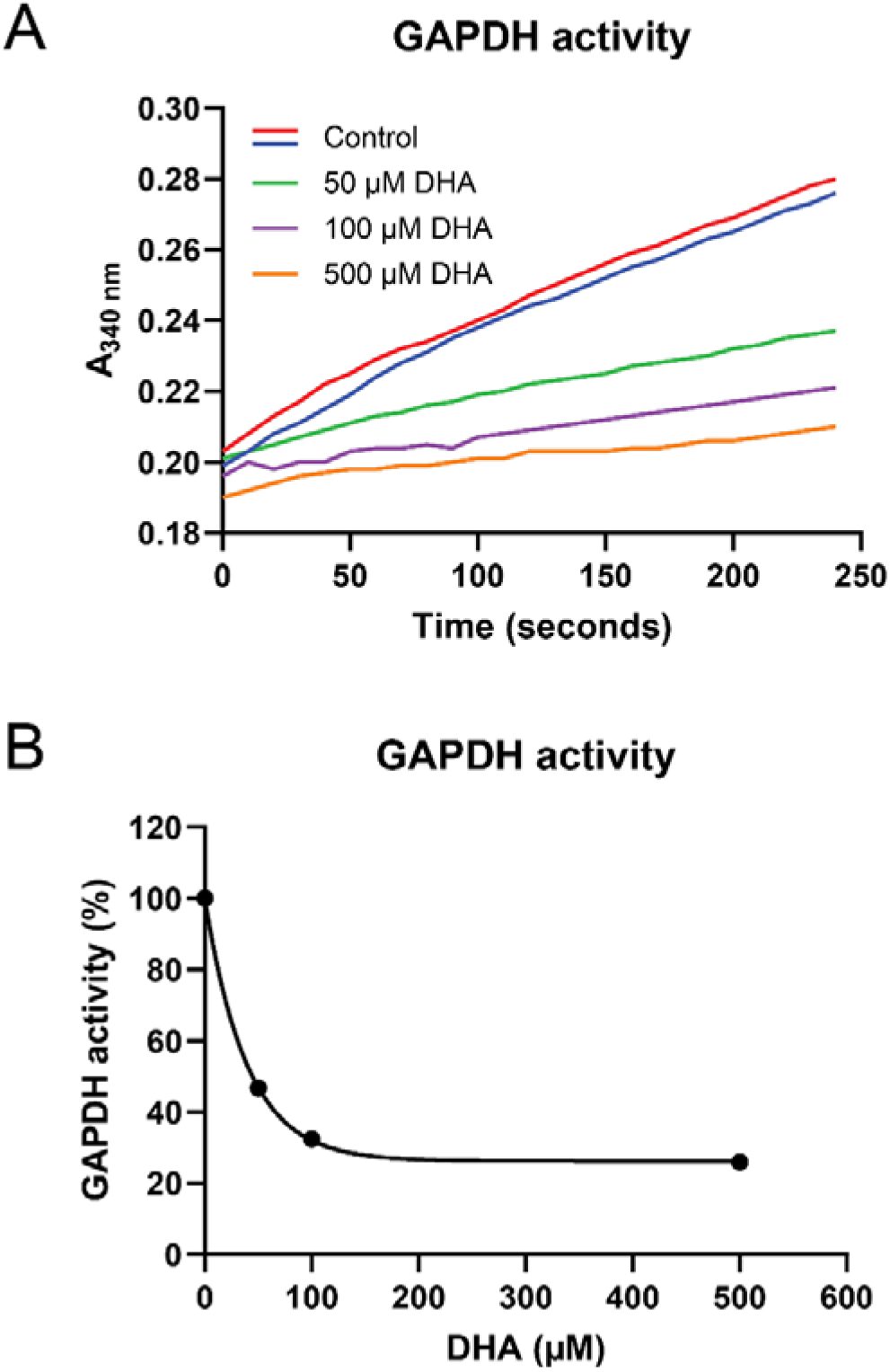
Inhibition of GAPDH by the products of ascorbate oxidation. Ascorbate was mixed with HOCl at a 2:1 molar ratio, followed by the addition of catalase (10 µg/ml). This mixture was then immediately added to GAPDH (see Methods). After 30 min, enzymatic activity was measured as outlined in Methods. The indicated concentrations of dehydroascorbic acid (DHA) are based on assumption that half of the ascorbate was oxidized by HOCl to form DHA. Control was GAPDH without treatment (A). Loss of GAPDH activity was dependent on DHA concentration (B).

#### 2.3. p16^INK4A^

The tumor suppressor p16^INK4A^ induces cell cycle arrest and senescence in response to oncogenic transformation. Oxidation of the single cysteine of p16^INK4A^ results in disulfide-dependent dimerization, which is a prerequisite of amyloid formation (37). LC/MS analysis showed that incubation of the protein with oxidized ascorbate for 60 min on two occasions did not result in modification of the p16^INK4A^ (Fig. 7).

**Figure 7.**
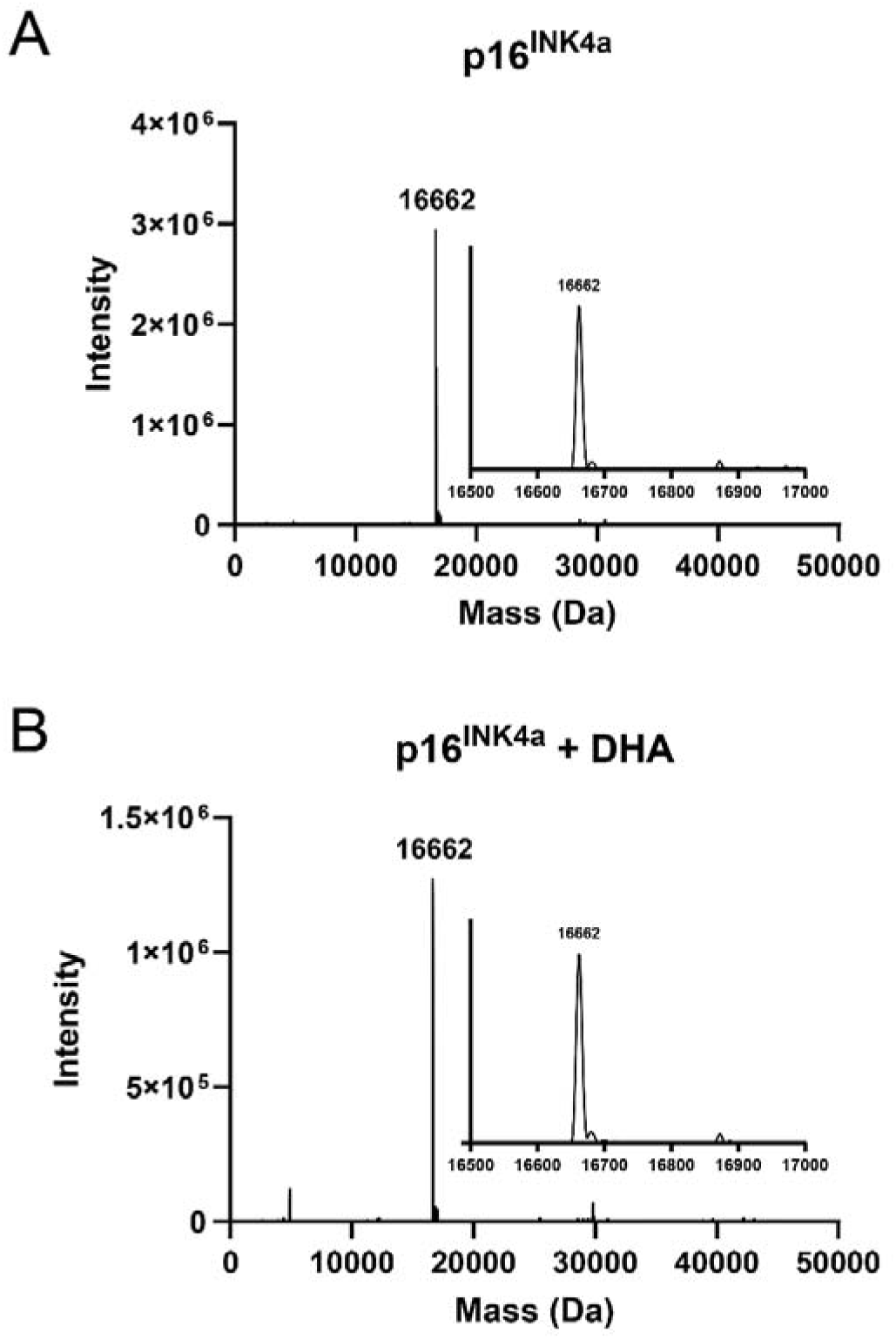
LC/MS analysis of the interaction of products of ascorbate oxidation with p16^INK4a.^ 10 µM p16^INK4a^ (A) was added to ascorbate+HOCl (2:1 molar ratio) mixed immediately before added to p16^INK4a^ (B) to give 500 µM DHA in reaction mixture. After 60 min incubation the mixture was applied on Micro Bio-Spin 6 columns (Bio-Rad) to remove unreacted low molecular mass molecules. No protein oxidation by DHA was detected on two occasions.

### 3. Effect of oxidized ascorbate on Jurkat cells

Jurkat cells, a leukemic human T lymphocyte cell line, was used to study the effect of oxidized ascorbate on cell morphology and viability. To do this, PI was added to label the DNA of dead cells and the number of cells was quantified by flow cytometry: the number of all cells in a given volume was counted and the percentage of PI-positive cells recorded. We observed a cytotoxic effect of 2.5 mM or 5 mM ascorbate, however in the presence of catalase ascorbate was well tolerated, with no visible changes in morphology or the number of viable cells, compared to controls (Fig. 8 and Fig. 9). In contrast, when ascorbate was oxidized with HOCl before addition to the cells, there were drastic changes in cell morphology, especially at 5 mM (Fig. 8). Notably, cell viability was greatly decreased in the presence of oxidized ascorbate, and this was only partly ameliorated by catalase (Fig. 9). Our data suggest that treatment with oxidized ascorbate slows down cell proliferation (Fig. 9B), but we’d need more data to confirm.

**Figure 8.**
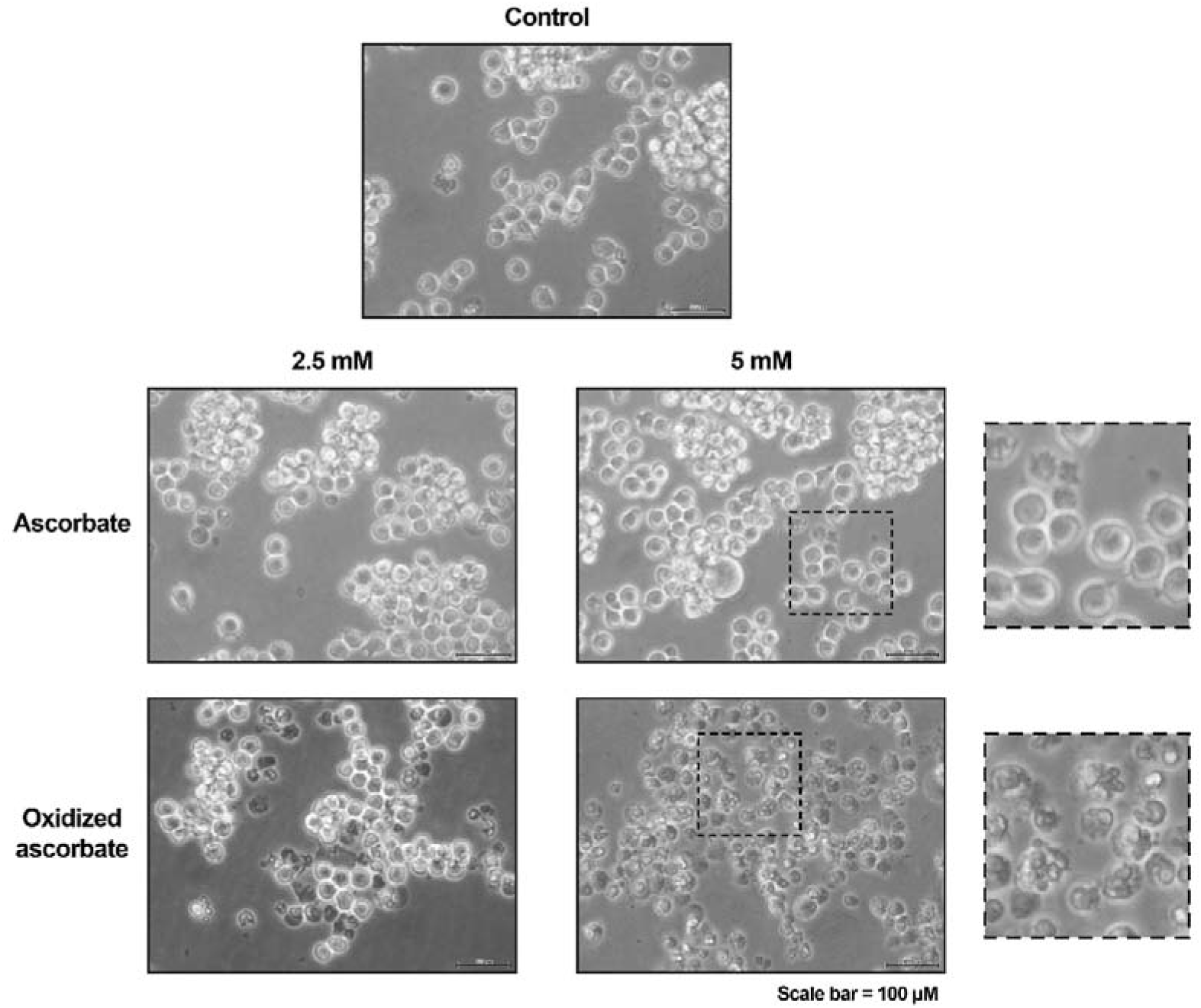
Morphological changes in Jurkat cells incubated in the presence of reduced or oxidized ascorbate. Cells were seeded at a density of 0.25 × 10^6^/ml before the addition of either ascorbate or oxidized ascorbate. To generate oxidized ascorbate, ascorbate (0.5 M) was mixed with HOCl (0.25 M), 1:1 (v:v) and added to cells in medium to achieve the required concentration. The stoichiometry of the reaction between ascorbate and HOCl is 1:1, indicating that oxidized ascorbate and the reduced form were present in equal proportions. Catalase, 20 µg/ml was also added to cells before ascorbate. After incubation for 24 h the cells were imaged using a 20x magnification objective, scale bar = 100 µm.

**Figure 9.**
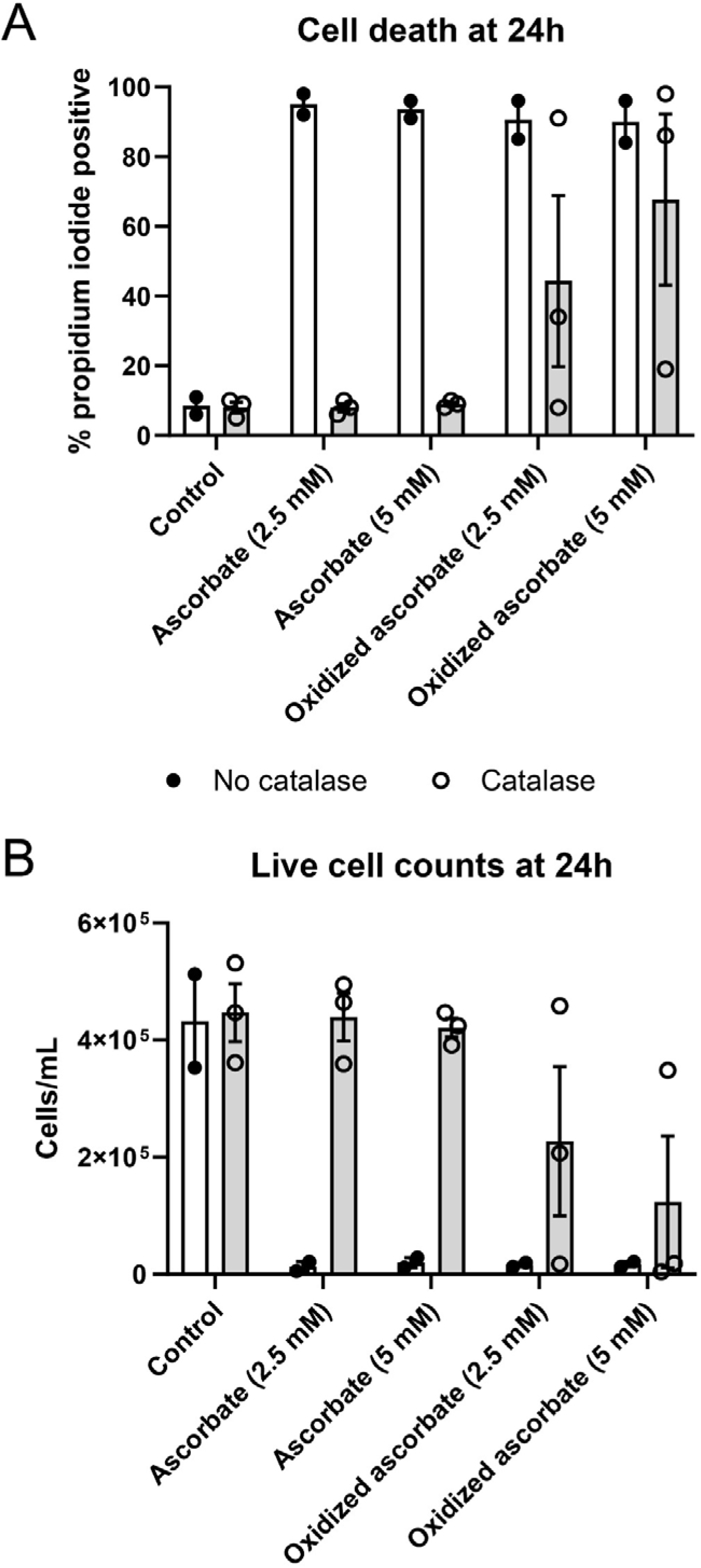
Viability of Jurkat cells after treatment with reduced or oxidized ascorbate for 24 h. To generate oxidized ascorbate, it (0.5 M) was mixed with HOCl (0.25 M), 1:1 (v:v) and added to cells in medium to achieve the required concentration. The stoichiometry of the reaction between ascorbate and HOCl is 1:1, indicating that oxidized ascorbate was present in equal proportion to the reduced form. Catalase (20 µg/ml) was added to cells before ascorbate. After 24 h cells were analysed by propidium iodide-dependent fluorescence using flow cytometry. (A) shows % dead cells and (B) the total number of viable cells. Opened symbols – cells with added catalase, closed symbols – cells without catalase added.

## DISCUSSION

Pharmacological treatments of cancer patients can be successful but are often confounded by i) a lack of selectivity, ii) the severity of side effects, iii) the development of tumour resistance to treatments, and/or iv) the appearance of secondary tumours due to carcinogenic effects on normal cells. In recent years there has been an interest in using high doses of vitamin C for cancer treatment. A growing body of evidence suggests that pharmacological vitamin C can be effective against tumour cells and generally no negative side effects have been reported. However, the anti-cancer mechanism of high doses of vitamin C is not fully understood and there is speculation that the full potential of vitamin C could be achieved in combination with other treatments. Currently, the anti-cancer mechanism of high doses of vitamin C is explained through pro-oxidant chemistry involving redox-active labile iron and H_2_O_2_ (7,8). Well-founded doubts remain however, that H_2_O_2_ generated by high doses of vitamin C could fully explain the anti-cancer effect, but no other feasible explanation has been suggested (9,10). We argue that anticancer effect of high doses of vitamin C is due to products of its oxidation. In this paper we observed oxidation of two thiol-dependent proteins, Prdx2 and GAPDH, which are recognized targets for anticancer treatment, by oxidized ascorbate. On the other hand, we report here that oxidized ascorbate had no effect on thiol of p16^INK4A^. It is well known that the reactivity of various thiols depends on the nature of the reagent, the protonation state of the thiol, and the surrounding amino acid composition (45, 47–50). Apparently, there is selectively in reactivity of oxidized species of ascorbate with thiols.

Activity of p16^INK4A^ depends on thiol oxidation, which is widely believed to be a common and important event in the development of cancer (37). The fact that p16^INK4A^ thiol is not susceptible to oxidation by the products of ascorbate oxidation suggests the selectivity of the anti-cancer effect of pharmacological vitamin C.

We have demonstrated that the viability of Jurkat cells was decreased in the presence of millimolar concentrations of ascorbate, and that catalase effectively protected cells. High doses of ascorbate are known to be toxic to cells in culture, and the protective effect of catalase appears to support the involvement of H_2_O_2_ as a culprit (51). Ascorbate is known to generate H_2_O_2_ in reaction with oxygen and also itself reacts with H_2_O_2_ (52, 53). By removing H_2_O_2_, catalase prevents the formation of DHA and thus inhibits the toxic effect. The toxicity of high doses of ascorbate could therefore be attributed to the generation of H_2_O_2_, followed by the formation of oxidized ascorbate, which was the bona fide cause of toxicity. Indeed, we observed cytotoxicity when a mixture of ascorbate and preformed oxidized ascorbate was added to cells in the presence of catalase. Observed toxicity to Jurkat cells warrants further study of the state of Prdx2 and GAPDH in cells to see whether they are inactivated after treatment with oxidized ascorbate.

Based on our results, we hypothesize that maintaining plasma vitamin C concentrations in the millimolar range through intravenous infusion, combined with MPO activation to facilitate vitamin C oxidation, could benefit anticancer therapy. The rationale is outlined below and summarized in Fig. 10, highlighting several potential mechanisms involved. Modification of Prdx2 and GAPDH may lead to disruptions in cell signalling and glucose metabolism, respectively. In addition, glucose metabolism can be suppressed through competition between glucose and DHA, the initial product of the two-electron oxidation of ascorbate, as both are taken up into cells by the GLUT family of transporters, which are overexpressed in tumors (54). The glucose concentration in blood is approximately 5 mM. Thus, by maintaining vitamin C at 10 mM or higher, one may induce competition for glucose transport into cancer cells. Combined with the inhibition of GAPDH by DHA, this approach could downregulate glucose metabolism. DHA can be effectively reduced by intracellular thiols, e.g. GSH (11), thioredoxin (Trx) (12) or Prdx (this paper). This could result in an elevated rate of thiol oxidation and increased oxidation of NADPH as it is consumed in reactions to reduce the disulfides formed. This might diminish the availability of Trx to ribonucleotide reductase, with a knock-on effect of lowering the production of deoxyribonucleoside precursors necessary for DNA synthesis, and resulting in decreased cell proliferation. Disulfidptosis is a recently identified form of programmed cell death induced by disulfide stress during glucose deprivation (55). Future studies may determine whether DHA can trigger this pathway.

**Fig. 10.**
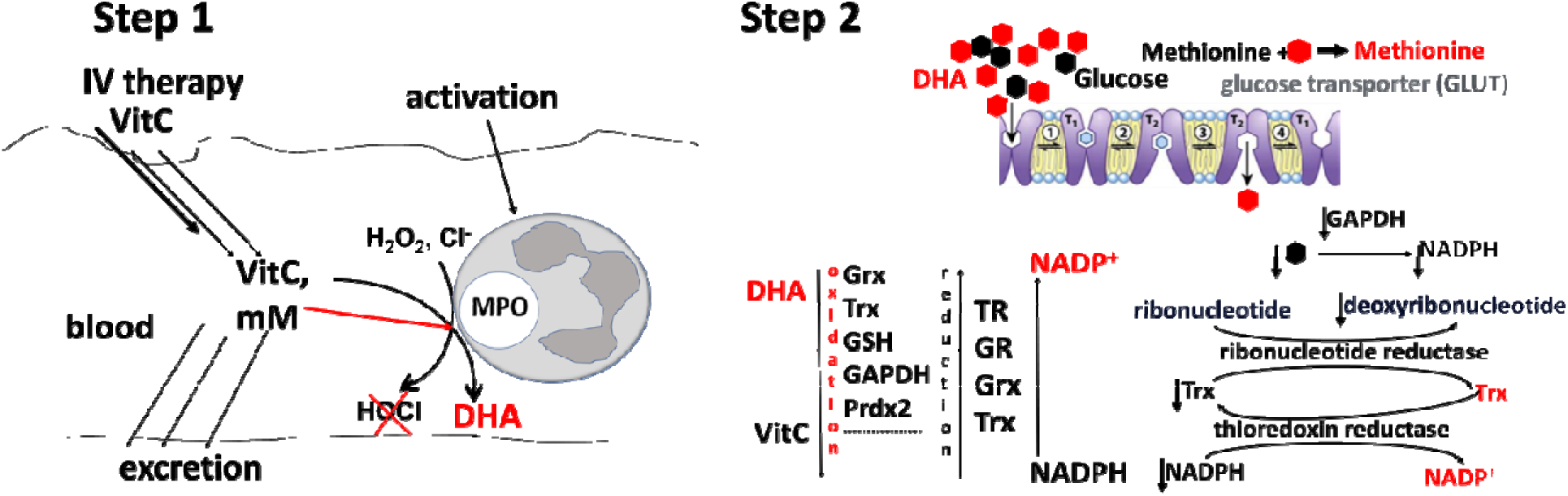
Administering a maintenance infusion of vitamin C to sustain a concentration in the millimolar range can effectively inhibit the generation of hypochlorous acid by activated myeloperoxidase (MPO) and promote the oxidation of vitamin C to its oxidized form, DHA (Step 1). DHA competes with glucose to enter cancer cells, leading to intracellular thiol oxidation and inactivation of critical proteins, subsequent downregulation of reducing power, and a depletion of precursors necessary for DNA synthesis. Oxidation of methionine by DHA may lead to methionine auxotrophy in cancer cells (Step 2).

Our observation of methionine oxidation by oxidized ascorbate suggests another potential pathway for the anticancer effects of high-dose vitamin C. Cancer cells are more sensitive to methionine restriction compared to normal cells. This phenomenon, known as methionine auxotrophy (56) —where cancer cells cannot grow without methionine while normal cells remain unaffected—may play a role in the anticancer effects of DHA.

It must be emphasized that the anticancer effect of vitamin C has only been observed with high doses and remains inconsistent. We believe that reported cases of successful anticancer therapy with vitamin C occurred when patients had inflammation, which assured high level of DHA.

Direct use of DHA for patient treatment would not be possible, due to its fast rate of decomposition (18,19). More promising would be treatment with vitamin C followed by its oxidation in situ. The oxidation could be performed by MPO from activated neutrophils: ascorbate could be effectively oxidized by HOCl, formed by Compound I of activated MPO (21) as well as directly by Compounds I, II, and III (22,57).

Here, we have demonstrated that millimolar levels of ascorbate effectively competed with physiological levels of chloride, leading to the generation of oxidized ascorbate and simultaneous inhibition of HOCl generation by MPO. For anticancer treatment, promoting vitamin C oxidation in the blood via activated MPO is appealing due to the high efficacy of this reaction.

The role of activated WBCs and MPO-generated HOCl in tumor initiation and progression is well documented (58–60). While activated WBCs and MPO can kill established cancer cells (61), the pro-carcinogenic effects of HOCl (62) and its involvement in various pathological conditions (63,64) pose significant challenges to its potential medical application.

The observed inhibition of HOCl generation by MPO, in the presence of millimolar ascorbate, may be a promising strategy for the treatment of various conditions in which the side effects of activated MPO are implicated (63,64). Although pharmacological companies are investing in the search for MPO inhibitors (65–67), a potent inhibitor of MPO, with demonstrated efficacy in human studies, has not yet been identified. Thus far the results of clinical studies using high doses of vitamin C in the treatment of sepsis have been controversial (68). The underlying problem which may inhibit a positive clinical outcome could lie in the fast clearance of vitamin C. With a half-life of approximately 2 hours, and the elimination of vitamin C from plasma following an exponential pattern, high plasma levels would be short-lived after a single intravenous injection (25). A continuous infusion aimed at maintaining blood ascorbate concentrations in the millimolar range could prove beneficial for inhibiting MPO activity in pathological conditions such as sepsis, where excessive neutrophil activity poses a concern. Concerns regarding oxidative stress in healthy tissues due to high DHA levels produced during the proposed treatment should be addressed in clinical trials. However, observations such as the high tolerance for vitamin C in patients (25), DHA’s selective inhibition of mitosis in L1210 leukemia cells but not in normal L929 cells (15), the lack of p16^INK4A^ oxidation by oxidized ascorbate, preferential uptake by cancer cells due to overexpression of GLUT transporters, and the higher rate of DNA synthesis in cancer cells suggest that this may not pose a significant obstacle. The observation that cancer cells, unlike healthy cells, do not exhibit biologically sufficient accumulation of the reduced form of vitamin C (69) suggests that high-dose vitamin C therapy may mitigate the risk of oxidative stress in normal tissues due to its potent antioxidant properties (70).

The combination of high-dose vitamin C with anticancer drugs has been proposed as a cancer treatment strategy (71). Indeed, such combinations have shown increased toxicity toward cultured cancer cells. For example, double-strand DNA breaks induced by the bleomycin–Fe(III) complex were significantly enhanced in cells loaded with DHA (69). This enhancement has been attributed to increased generation of reactive oxygen species due to iron cycling mediated by ascorbate, which is formed intracellularly through DHA reduction. Additionally, direct oxidation of thiol groups in DNA repair enzymes (72) by DHA may also contribute to this effect. Nevertheless, from a clinical perspective, such treatments may raise greater concerns regarding potential side effects.

## Acknowledgments

The authors express gratitude to Christine Winterbourn and Klaus Klarskov for valuable improvements to the manuscript. No funding support has been received from any grant funding body. No AI-assisted technologies has been used.

